# An agent-based model of the Foraging Ascomycete Hypothesis

**DOI:** 10.1101/197707

**Authors:** Daniel Thomas, Roo Vandegrift, Bitty Roy

## Abstract

Most trees host hundreds of species of fungi asymptomatically in their internal tissues, known collectively as fungal endophytes. The Foraging Ascomycete (FA) hypothesis proposes that some fungal endophytes inhabit the internal leaf tissue of forest trees in order to enhance dispersal to substrates on the forest floor, by using leaves as vectors and as refugia during periods of environmental stress. This dispersal strategy has been termed viaphytism. Following the FA hypothesis, many fungi may therefore be in continuous and cyclical flux between life stages as endophytes in the forest canopy and as wood-decomposing fungi on the forest floor. This cycle may represent a very common and previously-ignored process in the ecology of forests, with implications for forest health. The ecological consequences of the FA hypothesis are complex, so we constructed an agent-based model of the FA hypothesis. Our model is intended to serve as both an explicit conceptual explanation of the FA hypothesis, and as an exploration of the conditions in which a strategy of endophytism accompanied by leaf dispersal may be advantageous for fungi. In a scenario of a viaphytic fungal species on a model forest landscape, without fungal competitors, viaphytism is predicted to be a plausible alternative to dispersal to substrates by spores alone, allowing the fungus to persist reliably on the landscape. In a scenario that allows competition from aggressively dispersed non-viaphytic fungi, the model predicts some competitive benefits to fungal dispersal via leaves. However, these benefits are conditional, requiring sufficient retention through time of endophyte infections by host trees, and sufficient host trees on the landscape. In the model, loss of these fungal populations can result from increased local disturbances of forest canopy, and deforestation.

## Introduction

All large organisms are observed to host microbiomes (Rosenberg et al., 2010). Plants typically host very rich microbiomes, with both epiphytic (external) and endophytic (internal) fungi and bacteria present on and within all tissues (Rodriguez et al., 2009; Rosenblueth and Martínez-Romero, 2006; Vandenkoornhuyse et al., 2015). Additionally, a rich “endobiome” appears to be present at some level in all tissues of all plants. Fungal foliar endophytes (FFE) are an important subset of the plant microbiome, and have a long history of specialized study. We use here a common definition of endophytes: microorganisms that inhabit plant tissues for at least part of their life cycles, but do not cause disease symptoms, at least for an extended part of their existence within the host (Stone et al., 2000; Wilson, 1995). As with the microbiome at large, the FFE community is extremely diverse in most plants (Arnold et al., 2000; Arnold and Lutzoni, 2007), and FFE inhabit virtually all spaces of the leaf, especially in wet tropical ecosystems (Bayman et al., 1998; Gamboa et al., 2003; Gamboa and Bayman, 2001).

Interactions between host and FFE are complex, with relationships ranging from pathogenic to mutualistic (Carroll, 1988). Particular foliar endophytes have been observed to confer pathogen resistance (Arnold et al., 2003; Gazis and Chaverri, 2015; Mejía et al., 2008, 2007), drought resistance (Hamilton and Bauerle, 2012), other environmental stress resistance (Rodriguez and Redman, 2008), and defense against herbivory (Clay, 1988; Estrada et al., 2013; Sumarah et al., 2008). Even in the case of *mostly* commensal, neutral, or weakly antagonistic relationships with their host, the community of FFEs within a plant is hypothesized to act as an important modulator of disease, occupying space within plants that might otherwise be invaded by pathogens (Busby et al., 2016; Mejía et al., 2007). Microbial partners to plants may become more important in the current context of climate-change associated stresses (Busby et al., 2017; Woodward et al., 2012).

Nonetheless, the diversity and ubiquity of fungi living within leaves presents something of an evolutionary riddle. FFE species are often closely related to known plant pathogens, or are pathogens themselves when associated with a different host (Carroll, 1988; Freeman et al., 2001). Endophytic fungi have been found in the tissues of some of the earliest fossil examples of primitive land plants (Taylor et al., 2005; Taylor and Krings, 2005); for land plants, fungal endophytism may be as old as fungal disease. Further complicating matters, in contrast to the benefits for the host plant, benefits to fungal partners of an endophytic relationship are much less well understood. To be asymptomatic, endophytes must apparently restrict growth and sporulation while within their host (Stone, 1988), and there is evidence that the endophytic phase requires an array of unique metabolites (Carroll and Petrini, 1983; Kusari et al., 2012; Petrini et al., 1993; Schulz et al., 1999). An endophytic stage must supply some benefit to the fungal partner to outweigh these costs, or endophytism presents an evolutionary dead-end. The most common answer given to this riddle is the hypothesis that fungal endophytes are latent saprotrophs or pathogens (Carroll, 1988; Guerreiro et al., 2017; McMullin et al., 2019; Parfitt et al., 2010; Promputtha et al., 2007). Presumably, fungi living in leaves will have priority access to leaf tissues when host defenses weaken. Many foliar fungi do appear to be latent saprotrophs, meaning that they are very active in at least the early phases of leaf decomposition, consuming some of the more labile components of leaf tissue (Guerreiro et al., 2017; Osono, 2006; Stone, 1987). A portion of resident FFEs then sporulate on the fallen leaves they inhabit. Some of these FFEs are thought to overwinter on their decomposing leaves, sporulate in the spring, and return to the endophytic stage (Unterseher et al., 2013). However, many other fungi observed as FFEs have not been observed to sporulate (i.e., reproduce) on their leaves after senescence (Bayman et al., 1998; Lodge, 1997; Müller et al., 2001). These non-sporulating fungi must escape the leaf to another substrate after “robbing the bank”, or die without reproducing. This is particularly true in the face of intense competition from the diverse and aggressive, mostly basidiomycetous fungi that invade in the later stages of leaf decomposition (Voříšková and Baldrian, 2013).

Carroll (1999) proposed that some subset of endophytic fungi may utilize the endophytic phase to increase dispersal and colonization of woody substrates, a concept known as the **Foraging Ascomycete (FA) hypothesis** (Figure 1). He noted that leaves could act as excellent additional vectors for fungi residing within them, given a leaf’s ability to disperse relatively far from its tree (Ferrari and Sugita, 1996). Carroll also proposed that some endophytic fungi utilize the endophytic phase to not only cross spatial gaps to other substrates, but also temporal gaps between available substrate on the forest floor: fungi may take refuge in leaves to endure excessively dry or wet seasons that do not allow extensive decomposition of wood, or persist in leaves when woody substrates are not present. Thomas and Vandegrift et al. (2016) expanded this concept, noting that the humid microclimate provided by the shelter of a fallen leaf could protect endophytic hyphae as they grow out of a leaf, allowing endophytic fungi to escape from its host leaf to woody substrates, thus avoiding some of the uncertainty inherent in the germination phase of growth from spores. This form of dispersal also may coordinate dispersal times with events in which new substrates become more available, such as with storms that can both bring down new wood and leaves carrying fungi to the forest floor. At the time of the formulation of the original hypothesis by Carroll, few basidiomycete fungi had been observed as foliar endophytes (Carroll, 1988). Since this time, numerous basidiomycetous endophytes have been observed (for examples, see Arnold et al. (2007), Martin et al. (2015), and Thomas et al. (2019)), and may be candidates for the unique life history strategy suggested by this hypothesis.

**Figure 1.**
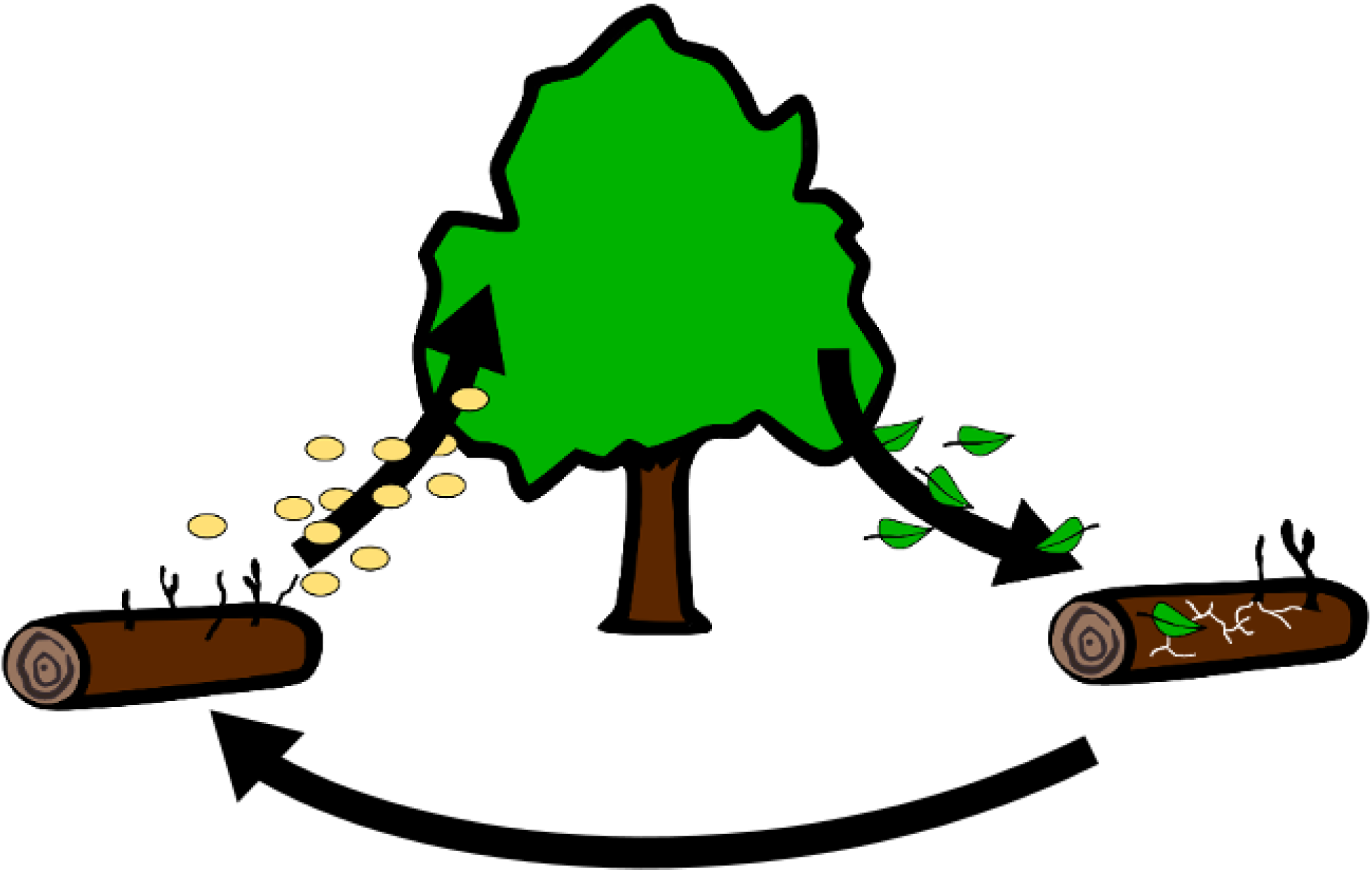
Visualization of the Foraging Ascomycete hypothesis, also known as “viaphytism”. Leaves are infected endophytically by spores, then act as dispersal vectors of fungi to wood on the forest floor. Fungi within leaves transfer to wood, decompose the woody substrate, and sporulate. Some of these spores germinate on leaves and colonize leaves again as endophytes.

The Foraging Ascomycete hypothesis centers on the competitive advantage given to fungi that can disperse by “hitchhiking” in leaves. Here, we use the term ***viaphytic***, recently proposed by Nelson et al. (2019), to describe endophytic fungi that transfer from endophytic infections in leaves to woody substrates, and thus are capable of using leaves as dispersal vectors. Fungi are increasingly shown to be dispersal limited, and increases in dispersal are therefore increasingly understood as vital for success in competition among fungi (Bruns, 2019). Viaphytism comes with the costs mentioned above that seem to make endophytism a curious bargain for fungi: production of enzymes for penetration of plant tissues and bypassing plant defenses, and subsequent suppressed metabolism and reproduction while occupying plant tissues until leaf senescence. However, for viaphytic fungi, we hypothesize that the costs of the endophytic phase “pay off” energetically and reproductively, because viaphytism externalizes some of the costs of dispersal to host plants, and ultimately results in a very competitive alternative dispersal strategy. Because of this alternative, leaf dispersal strategy, we hypothesize that the selective pressure on spore dispersal of viaphytic fungi is relaxed.

Conversely, non-viaphytes have evolved other ways to increase dispersal. There are numerous possible adaptations for directly increasing spore dispersal, such as by increasing spore quantities, optimizing spore or spore-bearing structure morphology, synchronizing spore releases for aerodynamic benefit, etc. (Fritz Joerg A. et al., 2013; Galante et al., 2011; Pringle et al., 2015; Roper et al., 2010).

The FA hypothesis is intriguing not only for the clues it may provide to the riddle of endophytes’ evolutionary history, but also for the large-scale ecological process it describes, which may be ongoing in forests everywhere around the world. Increasingly, fungi found as foliar endophytes are being located nearby as sexual-state fungi on woody substrates of the forest floor (Bills et al., 2012; Tanney et al., 2016; Thomas et al., 2016), very often *not* on the wood of their previous hosts (Lodge, 1997; Unterseher et al., 2013). The spores from these sexual fungi on the forest floor cause endophyte infections in nearby leaves, but this is not necessarily a one-way exchange. **The FA hypothesis proposes instead that these two great reservoirs of microbial diversity in a forest are in continuous and cyclical flux.** The scale of these processes should not be underestimated: in one Columbian cloud forest system, for example, 4.6 metric tons of leaves fall on a single hectare each year from the canopy, covering the forest floor several times over (Veneklaas, 1991). As all leaves examined so far have been shown to be largely saturated with FFE, this leaf fall represents countless potential transmissions of fungi to substrates on the ground (Nelson et al., 2019).

The process described by the FA hypothesis is not limited to wet tropical forests. The hypothesis was originally formulated to describe dispersal of FFE from conifer needles in the North American Pacific Northwest region (Carroll, 1999), and has been observed as possibly occurring in North American northeastern forests (Tanney et al., 2018). Given the potential commonness of the viaphytic cycle in forests, the importance of these fungi to the health of their host-trees (and therefore forest systems), and the role of these fungi in decomposing wood and leaf litter in forests, we argue that this ecological cycle has been both under-studied and under-appreciated.

To provide an in-depth theoretical explanation and exploration of the FA hypothesis as currently understood, we employed an Agent-Based Modeling (ABM) approach (Grimm et al., 2005). On a model landscape intended to simulate the site of research of a previous empirical study (Thomas and Vandegrift et al., 2016), we modeled a set of competition “experiments” between viaphytic fungi versus non-viaphytic fungi. We explored the ecological conditions in which viaphytism may endow significant advantage to viaphytic fungi in competition with aggressively dispersed non-viaphytic fungi. Additionally, given the current rapid rates of environmental change and deforestation throughout the tropics, we examined the resiliency of viaphytes when challenged by deforestation or increased loss of endophyte phase from the canopy.

## Methods

### Ecological system: fungi in the cloud forest

This model was inspired by an empirical study conducted by the authors in a primary cloud forest site at Reserva Los Cedros (www.reservaloscedros.org), in the northern Andes of Ecuador at 1300 m elevation (Thomas et al., 2016). Rainfall at the site can exceed 3 meters per year, and can occur daily throughout the year, often even during the dry season months of June through November. Forest structure is complex, with downed woody debris of all sizes, with an extremely high woody plant diversity of ∼290 tree species per hectare (Peck et al., 2011), and canopy gaps of varying sizes. Fungi, particularly fungi in the family Xylariaceae (Ascomycota, Xylariales), are often observed at this site both (1) as highly visible decomposer fungi, sporulating on woody debris, and (2) as microscopic endophytes within tree leaves overhead (Thomas et al., 2016), making it an obvious location for exploring the FA hypothesis.

The model landscape is intended to simulate a hectare of Andean cloud forest, after a recent storm or other gap-opening disturbance that has brought down more-than-usual woody debris. A disturbance of this small scale is reflective of actual canopy dynamics in many tropical forests (Espírito-Santo et al., 2014; Hunter et al., 2015; Nicholas V. L. Brokaw, 1982), and also represents the closest-possible “fresh start” scenario as can be imagined in the ancient ecosystem on which the model is based. Such an event may knock down fresh woody debris with lower fungal loads from the canopy to the forest floor, and blow new fungal spores into the site, or agitate the existing spore bank from the soil and debris. Additionally, leaf loss from the storm event and the new exposure due to the new gaps in the canopy at the site may stimulate the production of new, relatively sterile leaves at the site, ready for endophyte colonization.

### Software and computational resources

Model and scripts for simulations were coded in Python3 (www.python.org), using the *Mesa* agent-based model framework (Masad and Kazil, 2015). Additional core packages in the SciPy ecosystem were used to summarize and visualize results: *Matplotlib* (Hunter, 2007), *pandas* (McKinney, 2010), and *NumPy* (van der Walt et al., 2011). Simulations were carried out on the computing cluster in the center for Research Advanced Computing Services at the University of Oregon (https://hpcf.uoregon.edu/).

### Documentation

Much of technical detail of models are available in three online Jupyter notebooks. Online notebooks can be viewed by anyone using the links provided below, and are usually higher in quality than the supplementary pdfs, which are file conversions of the original jupyter notebooks. All code for processes and agents, and original jupyter notebook files used in the viewing links above are also available for download in public GitHub repository for this analysis (https://github.com/danchurch/FA_ABM).

Additional figures, additional results, and code for model runs from **parameter sweeps** of the scenarios explored below are available as a pdf (S1), or as an online notebook, viewable at <https://nbviewer.jupyter.org/github/danchurch/FA_ABM/blob/master/parameter_sweeps.ipynb>. Hereafter this is referred to as “notebook 1”.

Additional figures, discussion of calibration, and code for model runs of **dispersal parameters** in the models are available as a pdf (S2), or as an online notebook, viewable at: <https://nbviewer.jupyter.org/github/danchurch/FA_ABM/blob/master/calibratingDispersal.ipynb>. Hereafter this is referred to as “notebook 2”.

Additional figures, and code for calibration of **tree number and placement on the landscape**, including deforestation, are available as a pdf (S3), or as an online notebook, viewable at: <https://nbviewer.jupyter.org/github/danchurch/FA_ABM/blob/master/calibratingForest.ipynb>. Hereafter this is referred to as “notebook 3”.

Here we provide an informal summary description of our model and highlights of results of scenarios explored using the model. A formal Model ODD protocol description with further explanations of model concepts and assumptions is available in supplementary materials (S4).

### Model overview

The model is populated by three types of agents: fungal agents, tree agents, and wood agents. All agents exist on a flat, toroidal grid that is 100 cells in length and 100 cells in width, with each cell intended to simulate a square meter of forest. A general schematic of agents, processes, and scheduling is given in Figure 2.

**Figure 2.**
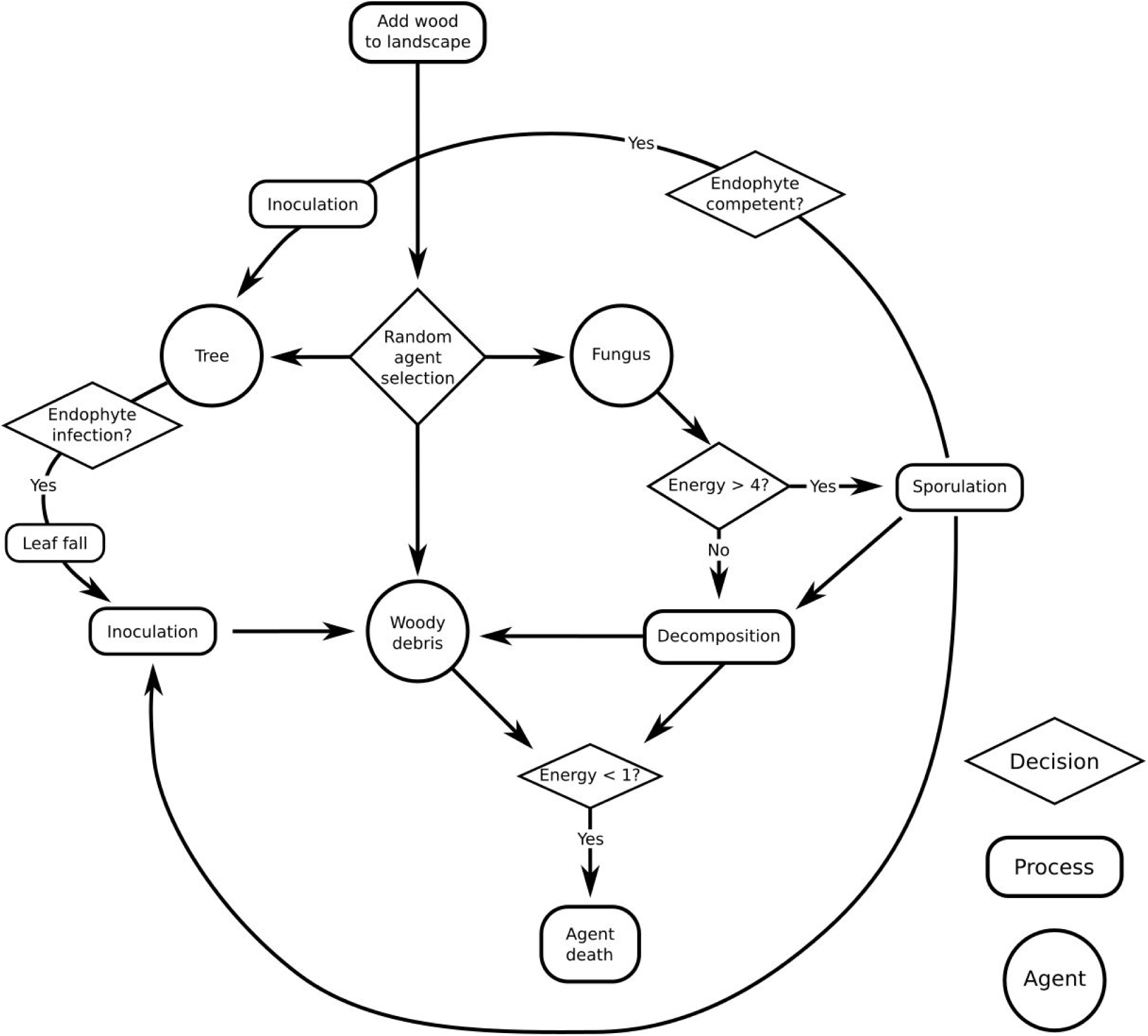
Schematic of processes possible during one timestep of the model.

#### Fungi

Fungal agents are the primary agents of the model. They initiate only on wood agents, and are capable of degrading the biomass of wood agents, and sporulating. They may be viaphytic, indicating that both tree agents and wood agents can be inoculated by their sporulation process, or they may be non-viaphytic, meaning that only woody agents are susceptible to inoculation by their sporulation. Some number of fungal agents are present on the landscape at the beginning of the model run, set by user. In all model runs presented here, one fungal agent is present at the beginning of the model run of either viaphyte, non-viaphyte, or one of each, depending on the scenario.

Fungal agents are initiated when a sporulation or leaf fall event successfully inoculates wood on the forest floor. Fungal agents then decompose the wood agent, storing 1 unit energy with each step as long as the wood agent continues to exist. 1 unit of biomass from woody substrate is removed for each unit of energy stored by the fungal agent. To model vegetative incompatibility systems in fungi, offspring fungal agents are generally considered vegetatively incompatible with their siblings and parents, maintaining independent accounts of energy and sporulation procceses. See ODD protocol (S4) for a detailed explanation of model assumptions concerning fungal nutrition/metabolism.

When they have sufficient energy, fungal agents undergo the process of sporulation. Cost of sporulation is set to 4 energy units, to simulate four 3-month timesteps, or yearly sporulation, if sufficient decomposition has occurred throughout the last several timesteps. Fungal agents reproduce and disperse through sporulation events and — if viaphytic — leaf fall events from infect tree agents. Neither spores nor leaves are modeled as *agents*; their behavior is modeled instead by the *processes* of sporulation and leaf fall. When a fungal agent reaches sufficient energy to sporulate, all wood agents on the landscape have a chance of being inoculated. If the sporulating fungal agent is viaphytic, all tree agents on the landscape with a negative infection status are also subject to inoculation. The probability of inoculation is a random variable that serves as a theta value for each bernoulli trial of inoculation, is defined as:

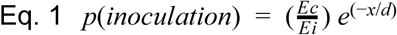

Where *x* is distance in meters between sporulating fungal agent and a susceptible wood or tree agent, E_c_ is currently energy/biomass of the wood agent, E_i_ is the initial biomass — thus making E_c_/E_i_ a handicap to further inoculation of a wood agent based on the state of decomposition of the wood agent — and *d* is the coefficient of dispersal that describes how aggressively dispersed a fungal agent is.

If the sporulating fungal agent an aggressive disperser, it will have a large *d* value, indicating that it has both a higher maximum radius of dispersal and a higher (“fatter”) local volume of spores (see Figure 3). The same process occurs for all tree agents on the landscape, if the sporulating fungal agent is viaphytic, without any decomposition handicap. If inoculation of a tree by spores is successful, the tree is infected by the endophyte and will disperse the fungus again to the forest floor via its leaves.

**Figure 3.**
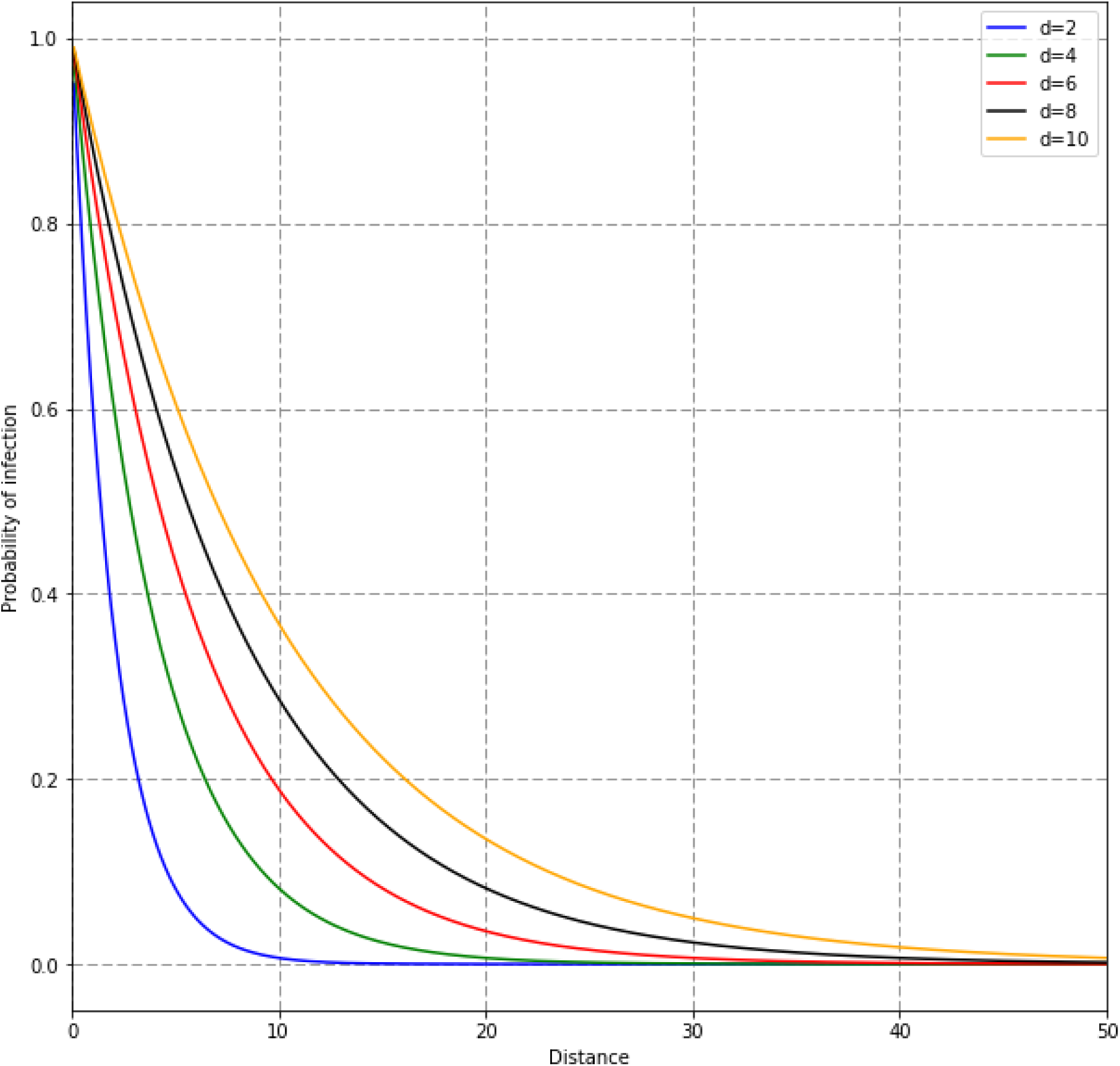
Distribution of probability for several different levels of dispersal ability according to our dispersal equation (Eq1). Distance units are meters. X-axis denotes the distance between a sporulating fungal agent or leaf-releasing tree agent, and a possible substrate. Spores of non-viaphytic fungi were calibrated from available literature and model runs to values from *d=*8 to *d*=10. Falling leaves were calibrated to *d=*4.

When a wood agent is no longer present, the fungal agent remains, respiring available energy reserves, as it searches for substrates until it dies, or until another wood agent falls onto the site. Vegetative (hyphal) discovery of new substrates is not modeled here. Dormant propagules are also not modeled.

#### Trees

Tree agents are located randomly throughout the landscape, according to a Thomas clustering process (Thomas, 1949), at a density similar to that found in global estimate of wet tropical forests (Crowther et al., 2015). Tree agents persist throughout a model run, unless deforested. Tree agents can be subject to deforestation, in a generally random (thinning) or random clustered (fragmented) model. See S3 or <u>jupyter notebook 3</u> on tree agent distributions and deforestation for more detail.

Similar to inoculations of wood agents by fungal agents, tree agents can also be inoculated during a sporulation event. Once a tree agent is successfully inoculated, its infection status changes from negative to positive. Endophytic infections are not directly considered fungal agents, instead these infections are tracked in the positive or negative infection status of a tree agent.

If a tree is successfully inoculated by fungal spores, they gain an endophyte infection. Each turn, an infected tree can stochastically lose its endophyte infection, at a probability set by the user. Tree agents shed leaves at regular intervals, capable of causing initiating viaphyte fungal agents if a tree agent has a positive infection status. Leaves are not agents; instead, leaf fall events are treated similarly to sporulations by fungal agents: they act as probabilistic dispersal events for transfer of endophytes to wood, using the same equation described above (equation 1). However, the dispersal coefficient *d* of leaves is much lower, to reflect the lower rate of leaves dispersed as compared to spores, and the lower maximum distance that leaves reach from the tree as compared to the farthest traveling spores. Additionally, leaf fall occurs every time step, unlike fungal sporulation. For more details about the calibration of leaf-based dispersal of fungi (viaphytism), see the jupyter notebook on dispersal calibration (S2 or <u>jupyter notebook 2</u>).

#### Wood

Wood agents appear on the landscape at every time step at a user-defined rate, with a random initial biomass. In addition, an initial population of wood agents is deposited at the beginning of every model run, at approximately twice the amount of wood brought in a normal season (three months), to simulate a recent decent disturbance. Wood agents have a single attribute, biomass. Biomass remains stable until wood agents are successfully inoculated with fungal agents either from a sporulation event or leaf fall event. Multiple fungal agents can inhabit a wood agent. Following inoculation, biomass is reduced by decomposition until zero, at a rate of 1 unit of biomass per fungal agent inhabiting the wood agent, until all biomass is respired away, at which point the wood agent is removed.

### Scenarios

Three scenarios were implemented, designed to answer the basic questions of the utility of a viaphytic dispersal strategy. All scenarios consisted of 100 model runs, where each run consisted of 50 timesteps, except for deforestation scenarios, in which each model run consisted of 100 timesteps. We created scenarios with the model intended to explore the following questions:

#### 1) When is viaphytism a beneficial dispersal strategy for fungi?

We hypothesize that the ability to disperse through leaves has reduced selection pressure on spore dispersal mechanisms of viaphytic fungi in evolutionary time. Thus viaphytes may disperse less aggressively through spores as compared to other competing fungal species, because their reproduction and dispersal is augmented by the additional dispersal strategy of using leaves as fungal vectors.

To quantify the extent to which viaphytism may represent a viable alternative to spore-dispersal, we compared the minimum threshold of spore dispersal ability necessary for a fungal agent to persist on the landscape indefinitely for both non-viaphytic (∧*d*_*nv*_) and viaphytic (∧*d*_v_) fungal agents. We propose that if *d*_v_ is significantly smaller than *d*_*nv*_, viaphytism may well provide dispersal benefits. In this scenario, viaphytic and non-viaphytic fungi were not competed directly against each other, but rather tested in isolation to determine their ability to survive in the default model conditions.

To find the minimal level of dispersal required for a non-viaphytic fungal agent to survive and flourish in the model landscape, we conducted a parameter sweep of the effect of spore dispersal abilities (“*d”*, equation 1) on the persistence of a non-viaphytic fungal agent in the default landscape. Resulting values for dispersal ability also were checked to conform to existing empirical data on fungal dispersal (Galante et al., 2011; Norros et al., 2012; Peay et al., 2012; Roper et al., 2010). At this stage we also tested the sensitivity of this “typical” non-viaphytic fungal agent type to initial reserves of woody debris on the landscape through a parameter sweep.

For viaphytic fungal agents, leaf fall process was calibrated according to data taken from the site (Thomas et al., 2016) and according to Ferrari et al. (1996). This leaf fall rate was held constant and assumed to be the rate at which viaphytic fungal agents could augment dispersal via leaves. Spore dispersal for a viaphytic fungal agent was then subject to a parameter sweep in a range of dispersal abilities as with the non-viaphytic fungal agent.

Refer to notebook 1 (S1 or <u>jupyter notebook 1</u>) for more details of the simulations to determine minimum threshold of spore dispersal ability necessary to persist on the landscape indefinitely for both non-viaphytic (∧*d*_*nv*_) and viaphytic (∧*d*_v_) fungal agents. Refer to notebook 2 (S2 or <u>jupyter notebook 2</u>) for comparisons of results to existing literature values.

#### 2) Is viaphytism a competitive dispersal strategy for fungi?

Viaphytic fungi are in competition with other fungal species for woody substrates. In the case of the Xylariaceous fungi that were the inspiration for this model, their competitors are the numerous white-rot basidiomycetous fungi that are rarely recovered as endophytes (Lodge, 1997; Lodge and Cantrell, 1995). We hypothesize that viaphytism may be an important trait for endophytic fungi to disperse to woody substrates, especially in the presence of intense competition from other fungi. In other words, we hypothesize that viaphytism may represent an evolutionary alternative to increasing spore dispersal ability.

A “typical” non-viaphytic fungal agent that could persist on our model landscape was defined above, in scenario 1. We further defined an “aggressive” non-viaphytic fungal agent as one that greatly exceeded the level of dispersal ability of this “typical” fungal agent. Thus, an aggressive non-viaphytic fungal agent was one with a dispersal coefficient *d* set significantly above the threshold needed to persist on the model landscape alone without competition (see results).

This “aggressive”, non-viaphytic fungal agent was then the benchmark competition, against which we competed viaphytic fungal agents of varying levels of dispersal *d* (equation 1). The dispersal ability of viaphytic fungal agents were incremented in different model runs until both populations consistently coexisted in perpetuity in an equilibrium, neither causing the extinction of the other.

Refer to notebook 1 (S1 or <u>jupyter notebook 1</u>) for more details of the simulations of competition between viaphytic and non-viaphytic fungal agents.

#### 3) Can viaphytes be lost?

We examined two ways in which viaphytic populations might be lost on a landscape: stochastic loss, and deforestation.

In natural settings, loss of endophytic infections could occur as a result of normal leaf senescence stochastically eliminating a small endophyte infection, or as defoliation events due to herbivory or disease, or in numerous other ways. In the current era of rapid environmental change, disturbances to forests such as disease and herbivory are predicted to increase (Ayres and Lombardero, 2000; Dale et al., 2001; Dukes et al., 2009; Seidl et al., 2017; Weed et al., 2013). Rather than anticipate each of all possible local disturbances which may cause host trees to shed FFE populations, local loss of endophyte infections was modeled as a stochastic process. During every time step, each endophyte-positive tree is at risk of losing its endophyte infection, at a user-set probability of loss. To test sensitivity of a fungal population to stochastic loss, we varied the probability of loss of endophyte infection from zero to extremely-probable (90% chance loss of endophytes per tree per time-step).

Deforestation was modeled in two ways, both intended to represent patterns of deforestation occurring in the region of our empirical study: (1) thinning, where trees are removed singly and throughout the landscape at an even rate, or (2) fragmentation, where contiguous blocks of forest are removed, each creating a gap with approximately 15 m radius on the landscape. The first attempts to emulate the results of selective logging, often in the form of “high-grading.” This is the most common form of deforestation at the site of research that inspired the model, and occurs continuously at unmeasured rates. The second is intended to model land-use conversions such as homesteading or conversion to pasture (Kettle and Koh, 2014), a generally common type of deforestation throughout the neotropics. Sensitivity of endophyte infections was detected by introducing deforestation into the landscape at the middle time point of 100-timestep model runs, at varying proportions of trees removed, under both styles of deforestation.

Refer to notebook 1 (S1 or <u>jupyter notebook 1</u>) for more details of the simulations of stochastic loss and deforestation and their effects on viaphytic fungi.

## Results

### Viaphytism as beneficial dispersal strategy

Viaphytism is a beneficial dispersal strategy as modeled here. A non-viaphyte fungal agent with a dispersal coefficient of *d*=8 was found to fit expectations from empirical data and to persist reliably on model landscapes (S1 or <u>jupyter notebook 1</u>). At this dispersal ability and higher, the fungal agents usually consumed the default amount of original woody debris on the landscape, then declined in population to match the rate of deposition of new substrates on the landscape. At lower than *d=*8 dispersal coefficient, populations often went to zero. These *d*<8 populations usually also had not finished decomposing existing wood deposits on the landscape by the end of the model run.

Without competition, fungi with an augmented *d*=10 dispersal ability only very rarely went extinct on the landscape. At this dispersal ability of *d*=10, non-viaphtyic fungal agents were able to fully colonize initial deposits of wood on the landscape, then maintain a steady population where all new woody debris was inoculated as it appeared on the landscape (Figure 4). We therefore considered this an “aggressive” non-viaphyte fungal agent, to be used in the following competition simulations with viaphytes.

**Figure 4.**
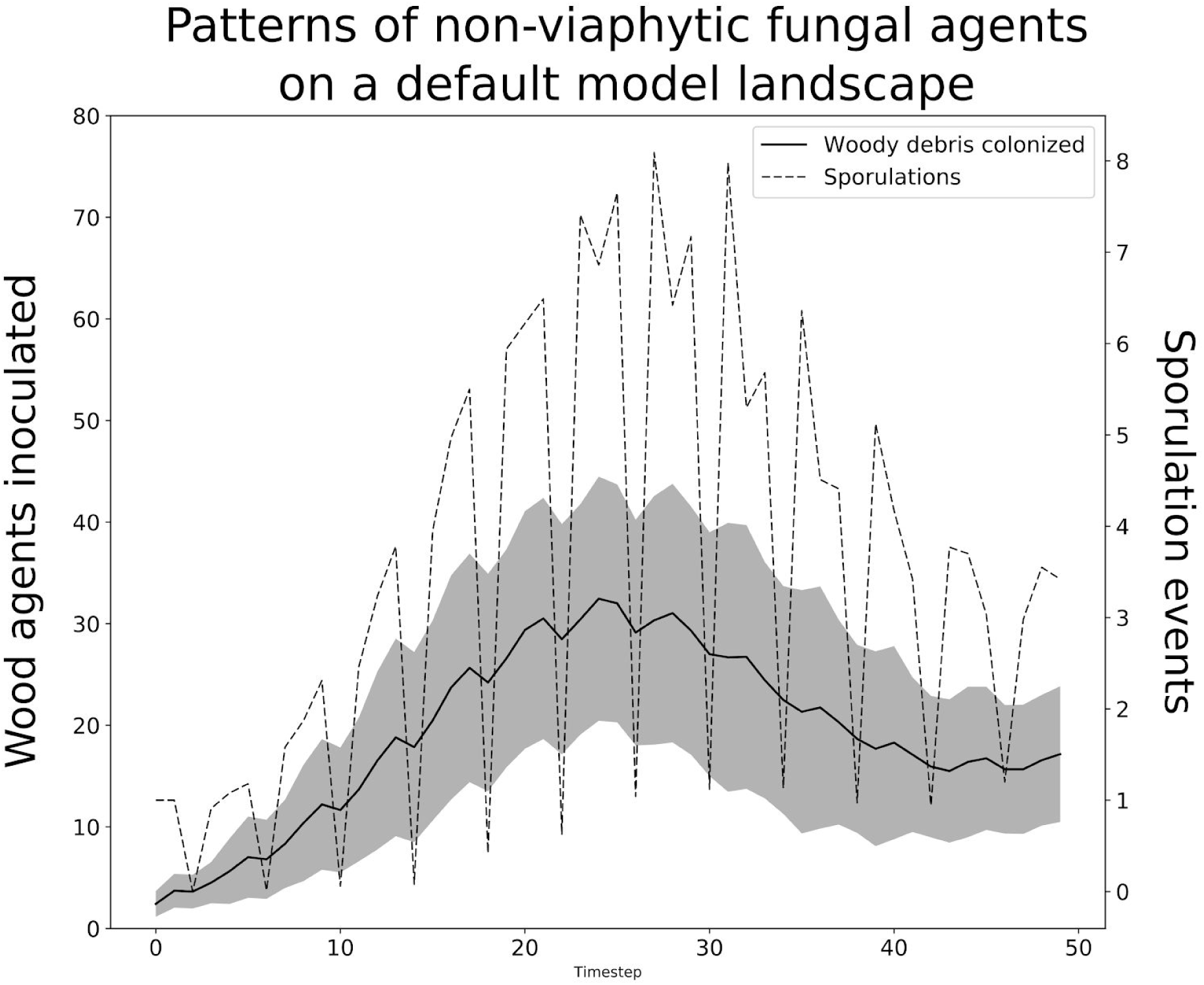
Behavior of an “aggressive” non-viaphytic fungal agents on the default model landscape. Error lines are one standard deviation from the mean.

Varying the amount of initial woody debris on the landscape had little effect on long-term behavior of the above “aggressive” non-viaphyte fungal agent type, as fungi typically consumed original woody debris in approximately the same about of time, though higher peak abundances of fungi were temporarily reached during the decomposition of this wood (Figure 5).

**Figure 5.**
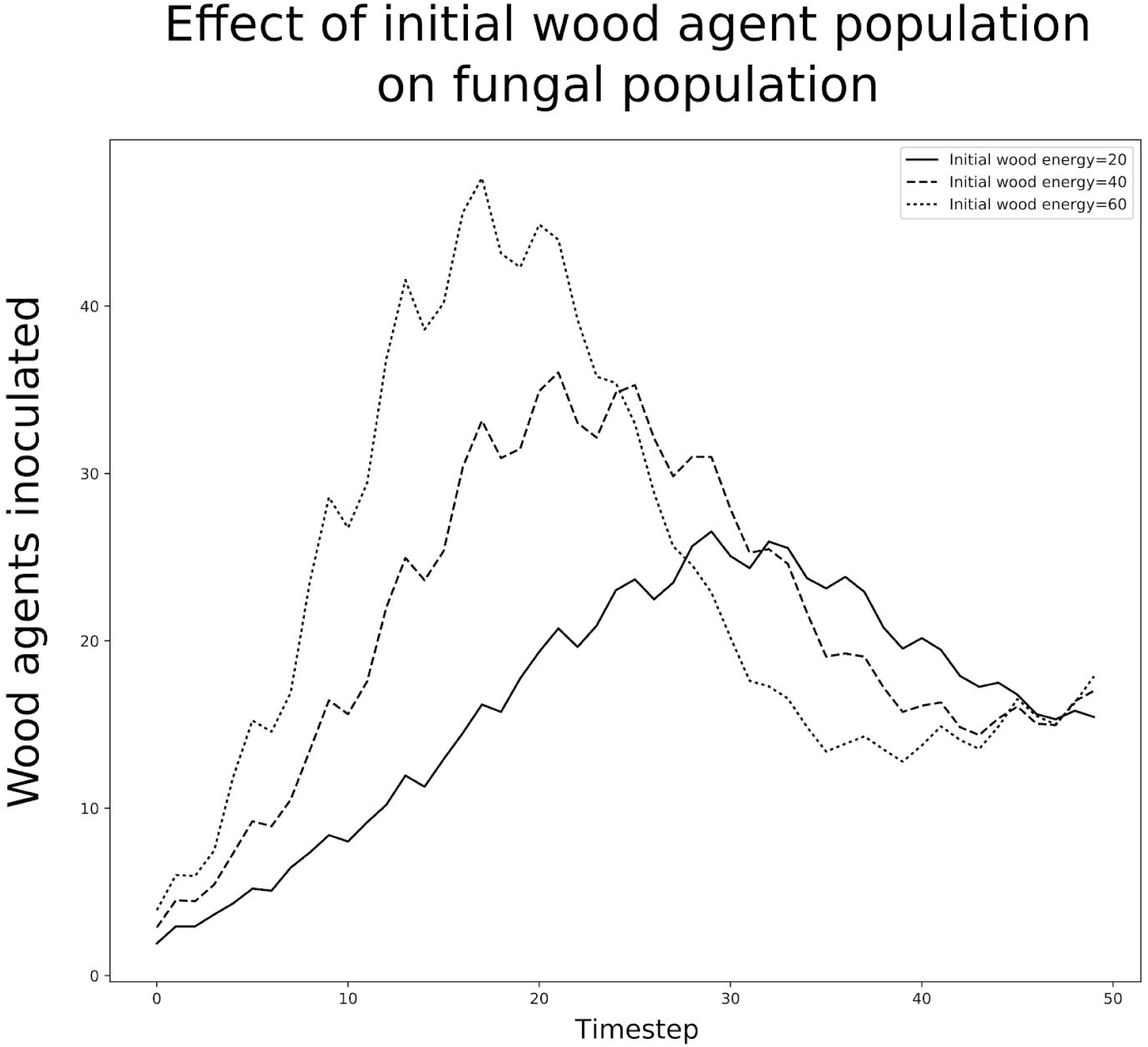
Response by fungi to varying amounts of initial substrate on the landscape. Within the range of values tested, varying the amount of initial amount of woody debris on the landscape changed only the early peak abundance of fungi on the landscape. Following this early peak, fungal populations that can persist equilibrate around the same population level, which is largely governed by the amount of new woody debris deposited on the landscape at each timestep.

Viaphytic fungal agents with a dispersal coefficient of *d*=2 persisted successfully very often on the model landscape, but did not entirely consume the reserves of woody debris on the landscape. (S1 or <u>jupyter notebook 1</u>). Viaphytic fungi survived in nearly all model runs and consumed original reserves of woody debris on the landscape before equilibrating to the rate of new wood deposition at *d*=3, just over a third of that necessary for non-viaphytic fungi to achieve the same equilibrium.

### Viaphytism as a competitive dispersal strategy

Viaphytic fungal agents with much lower spore-dispersal abilities compete successfully against aggressive non-viaphytic fungal agents (Figure 6). At *d*=2, viaphytic fungal agents can very often coexist on the landscape with aggressive non-viaphytic fungal agents. Below this dispersal level (*d*<2) for viaphytes, non-viaphyte fungal agents clearly outcompete the viaphytes, keeping infected trees and inoculated substrates by viaphytes to near zero levels. Viaphytic fungal agents with a dispersal coefficient of *d*=3 successfully and consistently outcompete the non-viaphytic fungal agents with a dispersal coefficient of *d*=10.

**Figure 6.**
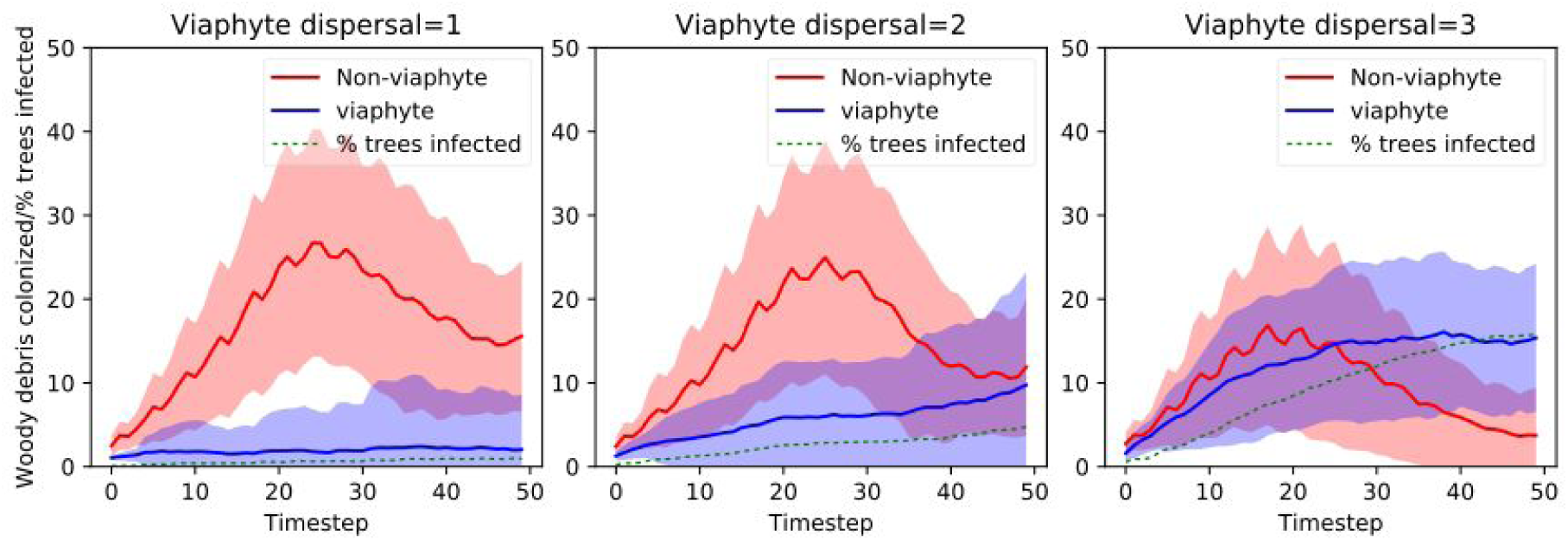
Competition of viaphytic fungi with various dispersal abilities against a model non-viaphytic fungal agent type. Error lines are one standard deviation from the mean.

### Loss of viaphytes from the landscape

#### Stochastic loss

All competitive benefits conferred by the endophytic phase to viaphytes as found in scenario 2 are dependent upon persistence of endophytes in the canopy. Under otherwise default settings, a probability of endophyte infection loss greater than 5% by tree agents caused loss of all competitive advantage by viaphytes against non-viaphyte fungal agents (Figure 7).

**Figure 7.**
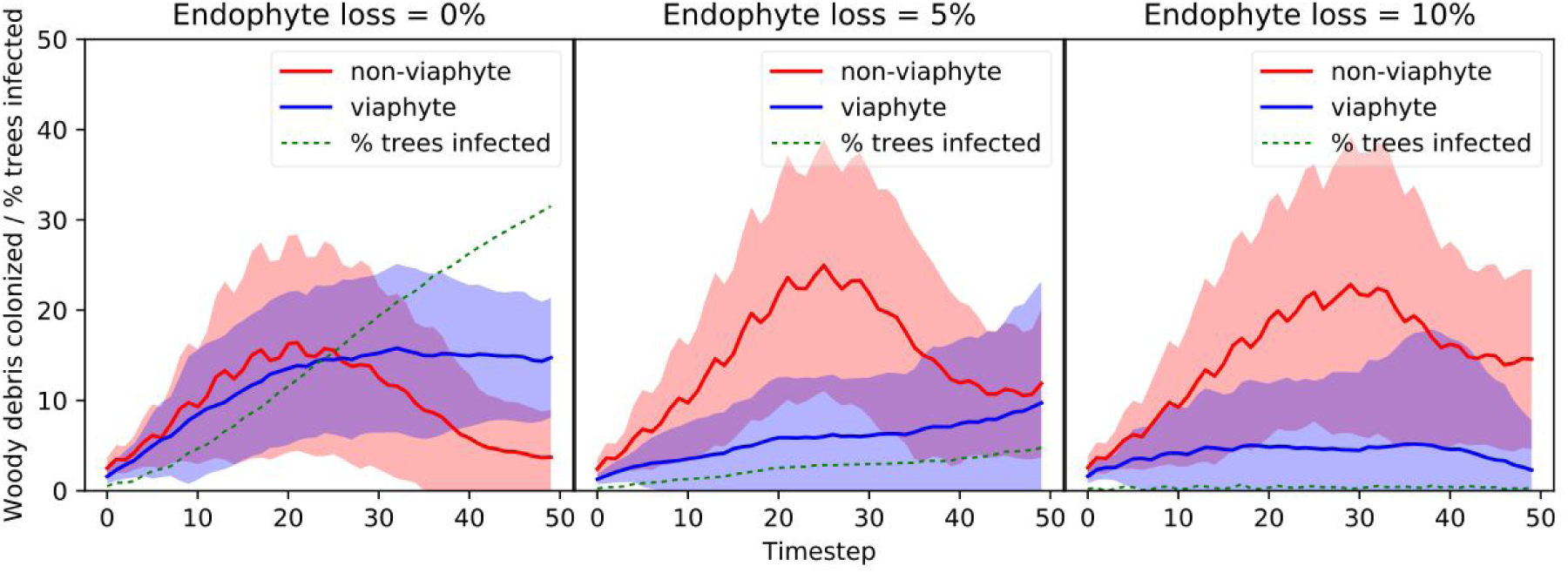
Effect of stochastic endophyte loss on viaphyte success. When tree agents have a higher than 5% chance of loss per time step, viaphytes show significant competitive disadvantage, often going extinct. Error lines are one standard deviation from the mean.

#### Deforestation

Consequences of removing trees depends on the intensity, timing, and spatial arrangements of the removal of trees. Without any cutting, model viaphytes show an increasingly stable presence on the landscape, as the reservoir of fungus in the canopy incrementally increases (Figure 8). Drastic thins (>70% of trees) reduce this stability, and hobble viaphytic fungi badly in competition with non-viaphytic on the landscape. Lighter thins (10-30%) appear to affect established populations of endophytes minimally. Fragmentation of forest showed similar effect as thinning simulations, on the spatial scale modeled here (S1, <u>jupyter notebook 3</u>).

**Figure 8.**
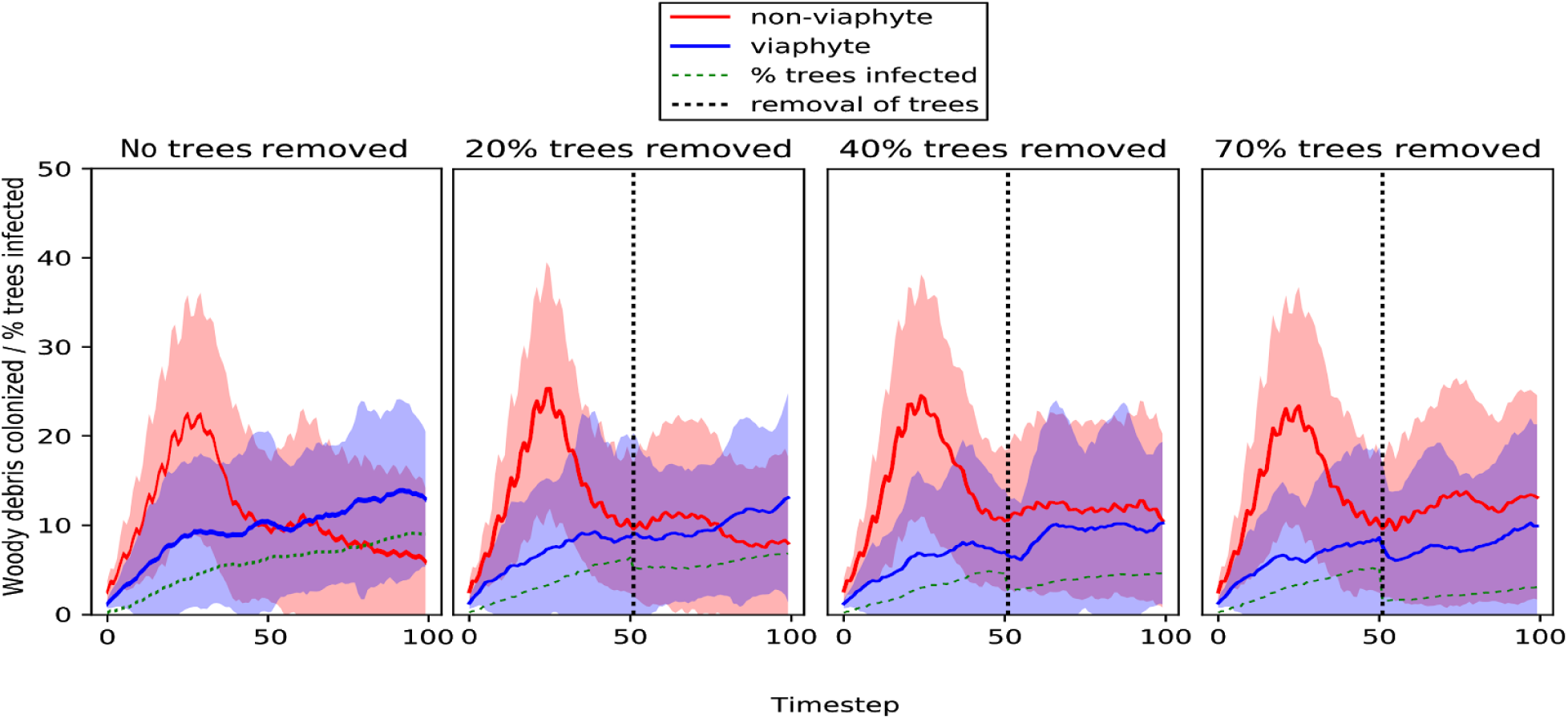
Effects of thinning trees from the landscape on viaphytic fungi, as they compete with non-viaphytic fungi. Trees were removed in a single event, at the halfway timestep of the model runs. Error lines are one standard deviation from the mean. More deforestation scenarios, including continuous thinning and small-scale fragmentation of forest, are detailed in S1 or <u>jupyter notebook 1</u>.

## Discussion

The model presented here predicts that some fungi may benefit from viaphytism, if endophytic infections can persist in the canopy for sufficient periods of time. As modeled here, the utilization of leaves as dispersal vectors and refugia in times of scarcity could allow a viaphytic fungus to persist and compete on a landscape of other, more aggressive non-viaphytic fungi.

This advantage is probably due, in large part, to some peculiarities inherent in reproduction by spores as a method of dispersal. When propagule dispersal is modeled as a negative exponential function of distance or related functions, as with the model presented here and elsewhere (Austerlitz et al., 2004; Galante et al., 2011; Levin et al., 2003; Rieux et al., 2014), abundance of spores and range of dispersal are coupled: most spores must fall locally in order for some small percent to reach farther distances. This sacrifice of numerous local spores to achieve longer range dispersal is often observed in nature, and is due to various mechanisms, such as with the synchronous ejection of spores from apothecia to reduce viscous drag from the air column (Roper et al., 2010), or the fluid dynamics of small particles in the air column generally (Levin et al., 2003).

In the model presented here, the effect created by the negative exponential model of spore deposition is highly visible during model runs, creating a positive feedback in colonization and consumption of substrates that is highly spatially and temporally autocorrelated. Once established, a fungal agent typically has high success in inoculating substrates locally, and thus it and its offspring typically rapidly consume locally available wood agents. If new substrate is not quickly found within dispersal limits, local extinction of fungal agents is very possible. Energy from wood agents are almost never limiting in the scenarios presented here, in a global sense - new wood agents are deposited at every timestep, at a rate far greater than is required by a fungal agent to survive and sporulate. Thus, it is not a global scarcity of substrate that drives populations of fungal agents extinct. Instead, it is the inability by fungal agents to disperse to other regions of the grid or the inability to persist locally while awaiting new wood agents to fall that drives populations of agents to zero. This “boom and bust” cycle of exponential growth and collapse is risky, and competition with other fungi accelerates this cycle in model simulations.

It is this risky aspect of spore dispersal that may make room for the viaphytic strategy. Viaphytic fungal agents, alternatively, may take refuge in — and augment dispersal with — an endophyte phase represented by a positive infection status in tree agents. Neither leaves nor spores of these endophytes are modeled as very widely dispersed, and they are often outcompeted locally for many timesteps in model simulations. Despite significantly handicapped spore-dispersal abilities, and despite only modest leaf-dispersal distances from infected tree agents, viaphytic fungal agents often displayed an incremental but reliable increase in population over on the grid over time, buffered somewhat from the boom-and-bust cycle experienced by non-viaphytic counterparts.

Though viaphytism as modeled here is a potentially beneficial strategy, its benefits may only exist under certain conditions. In the model, the potential dispersal benefits of viaphytism require sufficient time for endophytic infections to persist in the canopy. If trees do not maintain stable infections by endophytes in their leaves, viaphytism fails as a competitive strategy of dispersal. This is shown by the high sensitivity of viaphyte populations to greater than a 5% probability of loss from host tree per time step. It might be expected, therefore, that viaphytism is less common in seasonally deciduous forest systems, where yearly leaf fall presumably resets most endophyte populations to zero. In largely deciduous forests, we may instead more commonly see the yearly cycles of reinfection via spores from directly fallen leaves, as predicted by Unterseher et al (Unterseher et al., 2013). This sensitivity also suggests that the increasing disturbance regime predicted in many forests with climate change may impact fungi that rely on a viaphytic strategy for dispersal.

Viaphytism may also require sufficient numbers of trees on the landscape to succeed: in this model, if significant numbers of tree agents are removed from the landscape, a viaphytic life-stage ceases to be useful for overcoming dispersal limitation. Thus the FA hypothesis also points to more complex implications of deforestation. In such a system as we have modeled here, when a disturbance of the microbial community occurs on the forest floor (such as surface fire) or in the canopy (such as windthrow), recovery of microbial diversity and ecosystem services lost from one habitat may partially depend on the reserves of the other. Disrupting this cycling of microbes by deforestation in a modern, mechanized manner through tree-cutting may have unforeseen consequences to the secondary forest microbiome. Understanding modern deforestation in this way may help explain long-term differences observed in mycobiomes among different types of secondary forests and with nearby primary forests (Chaverri and Vilchez, 2006; Lodge, 1997). Removal of both reservoirs of fungal diversity (removal of forest canopies and resulting disturbance/exposure of soil) may mean greater loss of species, and slower regeneration of fungal diversity than smaller scale forms of timber harvesting and habitat conversion like slash-and-burn agriculture.

The work described herein was produced to be a theoretical explanation of the Foraging Ascomycete hypothesis. While this study defends the theoretical potential for this ecological process, further empirical exploration is now needed to gauge the actual importance of this process to forest ecosystem functioning and health. Such work is possible with laborious and invaluable studies such as those by Tanney et al (Tanney et al., 2018, 2016), which involve both classical, culture-based surveys of endophytes and intensive searches for cryptic macrofungi of the forest floor in the vicinity of the endophyte study. Further experimental exploration of the transmission of fungi from an endophytic state to a wood-decomposing state, such as done by Nelson (2019), is also needed.

## Author Contributions

D. Thomas, R. Vandegrift and B. A. Roy made wrote the text. D. Thomas wrote python code and ran simulations. B. A. Roy provided institutional support.

## Acknowledgements

We thank Dr. George Carroll for his interesting hypothesis, his inspiration as a mycologist and ecologist, and for all of his time given over the years in numerous theoretical discussions. We thank Research Advanced Computing Services at the University of Oregon for use of computing facilities necessary to complete all simulations. We also thank the anonymous reviewers for greatly improving the manuscript with suggestions and comments.

