## Supplementary material for "An agent-based model of the Foraging Ascomycete Hypothesis": (S4)

### Supplementary material 4: ODD (Overview, Design concepts, and Details) protocol summary

#### Purpose

The purpose of this model is to explore the feasibility of Viaphytism (Carroll 1999, Thomas and Vandgrift 2015, Nelson 2017), as part of a fungal life history and dispersal strategy. An ABM approach is used here to explore the possible advantages to fitness and dispersal conferred by viaphytism in fungi, by enacting competition-type scenarios among fungi with and without viaphytism.

#### Entities, state variables, and scales

Three agent types are placed on a spatial grid: trees, fungi, and wood.

Tree-agents represent individual adult trees with diameter-at-breast height greater than 10 cm. State variables of tree agents include position, leaf dispersal ability, state of endophyte infection (positive or not), and rate of endophyte loss. Tree-agents are treated as dimensionless, so that two tree agents are allowed to exist on adjacent cells, even though cells are intended to represent a square meter. Leaf dispersal ability is a positive integer, where larger values represent longer-range and more plentiful leaf deposition (see submodels). State of endophyte infection denotes whether a tree agent carries the endophyte stage of an endophyte-competent fungal agent in its leaves. Successful infection from fungal spores changes a tree agent's infection state to positive. Infections can be lost, and this loss is controlled by the endophyte-loss state variable, a number between 0 and 1, representing the probability that an infection is lost at each timestep.

A fungal agent represents a mycelium, resulting from a single reproductive event, either a spore- or leaf-vectored inoculation of wood. State variables of fungi include: position, spore dispersal ability, stored energy (biomass), and viaphyte-competence. Like leaf dispersal with tree agents, spore dispersal ability is a positive integer, with larger values representing longer-range and more plentiful spore deposition across the landscape when sporulation occurs (see submodels). Energy is representative of biomass and potential energy gain from decomposition of wood agent. Sufficient energy stores allows for a sporulation event. Viaphyte-competence denotes the ability of a foliar fungal endophyte to transfer from fallen leaves to woody substrates. In terms of the model, viaphytic competence indicates whether a fungal agent can change the endophyte infection status of a tree agent during a sporulation event, and then disperse through leaves to inoculate wood agents.

Wood agents represent the biomass deposited on the forest floor from the canopy. State variables of wood are position and stored energy (biomass). New wood are given a starting amount of energy, and this wood biomass is converted incrementally to fungal biomass if fungi are present in the cell.

Grid cells are not given attributes, except for the agents they hold, and their location, in the form of x and y coordinates. For all the scenarios examined here, the grid spans one square hectare (100m by 100m), wherein each grid cell represents one square meter. Grid is flat, and toroidal. Agents of all types can occur at all grids, though fungi will not persist for long periods without wood agents also present because of energy constraints.

Model-wide, environmental state variables include the rate of deposition of new wood agent, number and spatial clustering parameters. Trees can be removed at any time during a simulation to model effects of deforestation.

#### Process overview and scheduling

Time steps begin with the placement of new wood agents on the landscape. Following this, agents are chosen randomly to act, regardless of type. See figure 1 for a summary schematic of model processes for one time step. All random numbers for scheduling, and for all submodels are drawn from the uniform distribution, with minimum and maximum values defined by the nature of the process and number of agents involved.

Fungal agents begin with a test of their biomass (energy) reserves. If energy is high enough, sporulation occurs, possibly instantiating new fungal agents on wood agents. If the sporulating fungal agent is viaphytic, the spores can also change the endophyte infection status of tree agents on the landscape to positive. Sporulation results in a loss of energy for the parent fungal agent. Following this, fungal agents decompose the wood agents available in their grid cell, resulting in a gain of energy for each fungal agent present and a loss of stored energy in the wood agent. If the wood agent at a grid cell has died, fungal agents continue to respire, subtracting from their energy each turn until they have energy $< 1$, upon which they die.

Tree agents begin by dropping leaves. If a tree agent has a positive endophyte infection state, these leaves disperse to the landscape and can inoculate wood agents, instantiating a new fungal agent. Trees can also be removed from the landscape, which if requested occurs at the very beginning of a step, before deposition of wood agents.

Wood agents are placed at the beginning of each time step, in multiple random locations at the start of each step. The exact number of wood agents laid down each step is random, but the energy in each varies and the total energy represented by all the new wood agents will approximately equal the New Wood Energy state variable set by model user. Wood agents then test their biomass (energy) state variable: when energy $< 1$, the agent is removed from the landscape.

After all agents present on the landscape act, data collection takes place, and the time step is complete. Under model default settings, each time step is intended to represent approximately 3 months.

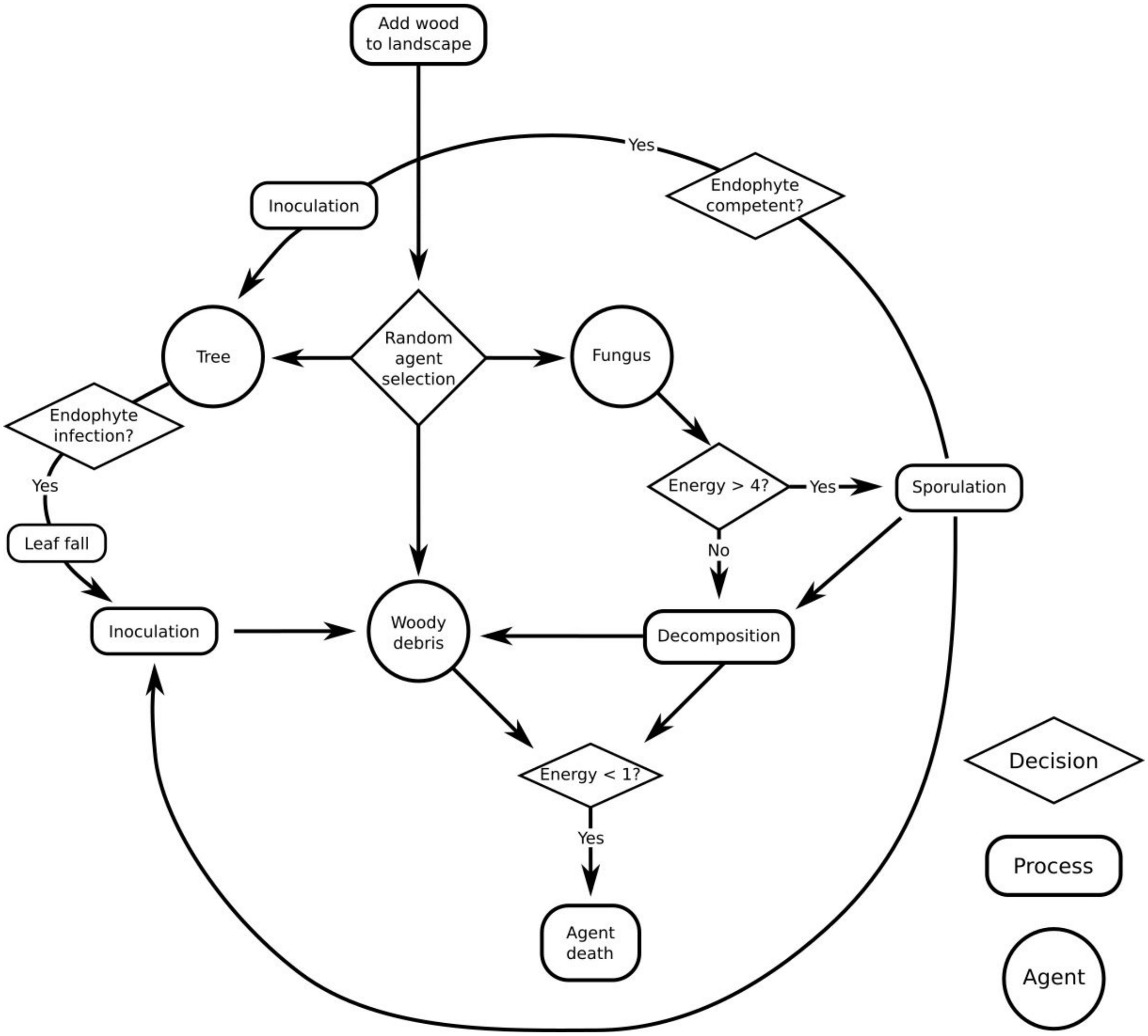


Figure 1. Schematic of processes possible during one timestep of the model. [For a higher resolution image click here](#).

#### Design concepts

##### Basic principles

This ABM is primarily a model of dispersal and competition among fungi. Patterns of spore dispersal at various scales are measured in Galante et al (2011), Norris et al. (2012), Peay et al

(2012), and others. These studies show that the negative exponential family of functions can be parameterized to fit abundances and probabilities of spore-dispersal in nature. Leaf fall has also been shown to be well described by exponential decay functions (Ferrari and Sugita 1996). These well-established patterns of dispersal serve as first principles in this model, guiding the behavior of both tree and fungal agents.

However, the purpose of the model is to explore the hypothesis that some fungi utilize an endophytic life stage to enhance dispersal and to persist on the landscape during times of scarcity, intense competition, or environmental stress (Carroll 1999, Thomas and Vandegrift 2015). This viaphyte life history strategy, where some fungi alternate endophytic and free-living phases, is a basic principle of the model, and the focus of the simulations presented below.

##### Emergence

Since agent learning and evolution are not allowed in the model, possibilities for emergent properties and patterns are limited with this model.

##### Adaptation, Objectives, Learning, and Prediction

Fungal agents seek reproductive success, which can be measured either by number of substrates occupied or sporulation events. However, fungal agents are not given the ability to modify their behaviors to increase fitness. As such, they do not take any measure of success, memory of past events, or predictions of future conditions, into account during their actions.

##### Sensing

Fungal agents' decisions are based primarily on internal sensing of biomass (stored energy) to decide when to initiate sporulation and external sensing of distance wood agents and tree agents when sporulating, to determine the probability of infection. Inoculation of wood agents by endophyte infected tree agents also senses the distance to wood agents to calculate probabilities of infection.

##### Interaction

Interactions among fungi are indirectly competitive, mediated through wood debris agents. Wood agents are consumed by fungal agents as a source of energy, and the presence of existing fungal agents associated with a wood agent reduces the likelihood of establishing new fungal agents on a wood agent. This is enforced by the $E_c/E_i$ ratio that is a coefficient in our equation for calculating probability of inoculation success by spores or leaves onto wood. This is discussed in the wood decomposition submodel.

##### Stochasticity

Several stochastic processes are used in the model to emulate the variable environment of forest ecosystems. Amount of wood deposition per step, number of successes in sporulation/inoculation, initial placement of trees and wood agents, and methods of tree agent selection in deforestation all involve stochastic selections of agents and locations. These are described in the submodels.

##### Observation

At the end of each step in the model the following are recorded: total numbers of fungal agents, wood agents occupied by fungal agents of both endophyte-competent and non-competent fungi, total sporulation events by both types of fungi, percent of tree agents infected by endophytes, and for deforestation scenarios, total number of tree agents on the landscape.

#### Initilization

Model default density of ~600 tree agents in a 100 cell × 100 cell grid are intended to approximate a 1 ha plot in wet tropical forests (Crowther et al 2015). Initial conditions of the model are intended to emulate a recent small disturbance in a forest landscape, where a larger than usual amount of uncolonized woody debris has been randomly deposited. Unless otherwise specified, all model runs begin with one fungal agent of each type, randomly associated with a wood agent. These initial fungal agents are assumed to have established themselves and begin the model with a starting energy sufficient to sporulate 2 or three times. Endophytism in the model can be disabled, allowing competition experiments between two non-viaphytic fungi. Dispersal coefficients are assigned to both types of fungal agent, and to tree agents for dropping leaves, though this last setting is typically held at a default value from leaf fall data (see submodels). Default initial Wood agents have a total biomass/energy of 30 (this can be changed by the user). Rate of new wood agent deposition on the landscape can also be set prior to initialization, though this was typically held a default value found to allow aggressive, non-viaphytic fungi to persist on the landscape. Initialization states are intended vary among model runs, to explore the benefits and limits of a viaphyte-style life history strategy.

#### Input

Deforestation scenarios require time-series input data, in the form of timing, intensity, and spatial nature of tree agent removal. Otherwise the model does not require input data.

#### Submodels

Submodels are listed in figure 1 schematically as processes. In addition, we describe procedures for initial placement of tree agents, and two deforestation submodels.

##### Wood deposition

Wood deposition (fig. 2) is given total energy budget per timestep (A), that is defined by the user/defaults before initiating a model run. To simulate the variety of sizes of woody debris that occur in forest settings, however, each new wood agent (W) is given variable (random) initial energy (e), taken from the iteratively smaller range of energy remaining. As agents are added, a tally of energy used ("a") is maintained. This tally "a" will ultimately approximately equal the wood deposition rate given by the user/default, and the submodel exits.

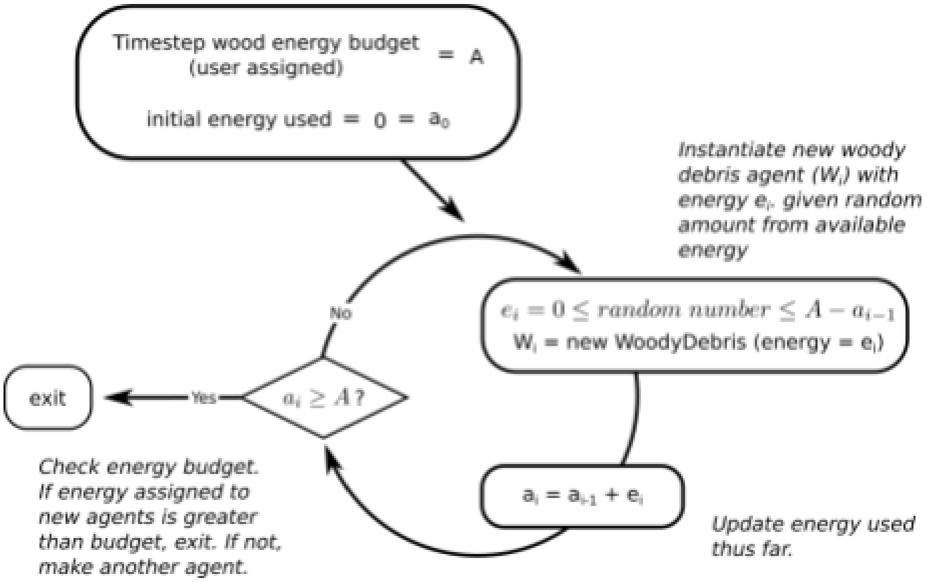

Figure 2. Wood deposition submodel. For a higher resolution image click here.

#### Sporulation and inoculation

Whenever a fungal agent sporulates, or an endophyte-infected tree agent releases leaves, every uninfected tree agent or wood agent is then subject to a Bernoulli trial, with a theta that is calculated from distance between targeted agent, and the agent that is the source of inoculum (sporulating fungal agent or tree agent). Calculation of probability of infection of a wood agent or tree agent from spores is an exponential decay function of distance ("x") from self (fungal agent), multiplied by a dispersal ability coefficient ("*d*") assigned by the user (figure 3). Viaphytic and non-viaphytic fungi can be - and usually are - assigned distinct dispersal abilities. Probability of Inoculation of wood agent is furthered multiplied by the fraction of current, remaining energy ("$E_c$") over starting energy ("$E_i$"), to give a handicap to colonization of wood agents by new fungi, if the wood is already inhabited by other fungi. For bernoulli trials, random numbers are drawn from the uniform distribution, between 0 and 1.

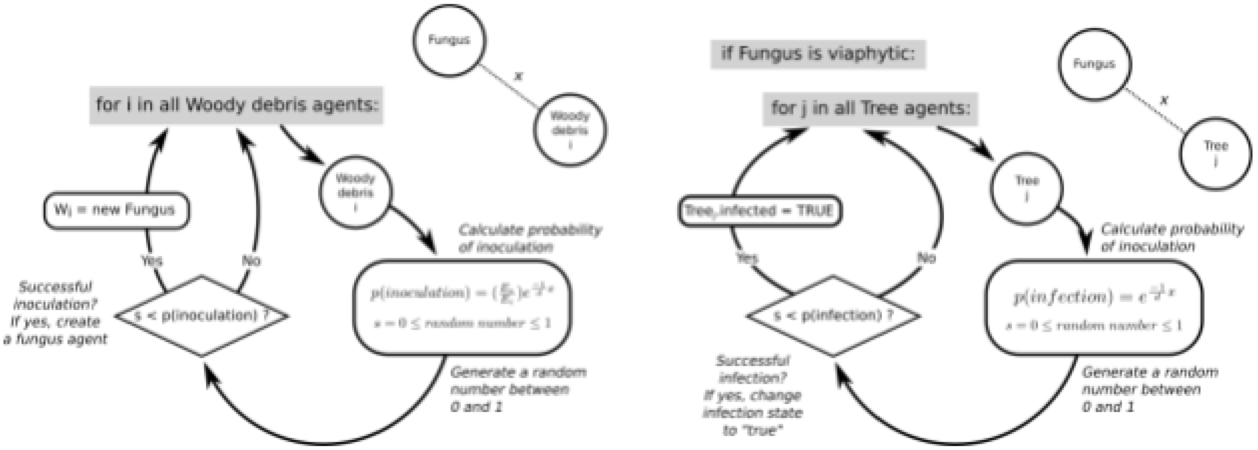


Figure 3. Sporulation submodel, for both wood agent inoculation and endophyte infection of tree agents.
For a higher resolution image click here.

#### Leaf fall and leaf-vectored wood inoculation

Leaf fall is treated similarly to sporulation (fig. 3), except that it occurs at every time step, as an action of all Tree agents, without any energy budgeting. For the purposes of this study, leaf fall for all tree agents was calibrated at d=4. The equation for determining the probability of inoculation of a Wood agent is identical:

$$p(inoculation) = (\frac{E_c}{E_i})e^{\frac{-1}{d}x}$$

Where "$E_c$" and "$E_i$" are current and initial energy, "x" is the distance between Tree agent and Wood agent, and "d" is the dispersal ability coefficient for tree agents, in the author scenarios these are held at d=4.

#### Decomposition

Decomposition is modeled here as a simple one-way transferral of energy from Wood agents to their associated Fungal agents. Every time-step, each Fungal agent on a grid cell with a Wood agent gains one energy, and causes the wood agent to lose one energy. Thus, a cell with numerous Fungal agents will show rapid decomposition of the resident wood agent, and becomes increasingly difficult to for new Fungal agents to access.  After a wood agent drops below one energy in a turn, it is removed from the model. Fungal agents will then lose one stored energy unit at a rate of one per step until dropping below one unit of energy, then removal from the model, unless a new wood agent is placed on the cell.

Biological rationale of decomposition process:

Units of woody agent biomass and fungal energy units are highly abstracted. Their exchange is a large simplification of natural processes, *not* intended to strictly model balances of fungal respiration, biomass gains in fungi, and biomass loss by woody substrates. Modeling the separate costs of respiration and growth (biomass accumulation) with fungi in nature are extremely difficult, precisely because they are not separable. This is true because hyphal fungi in nature are osmotrophic feeders that are thought to only rarely show true chemotrophy (Moore, 1998), and that most of their metabolic activity is to be found near active hyphal tips (Moore et al., 2011). This means that metabolic activity, including both growth and respiration, increases non-linearly when food is available, as fungi in nature must branch extensively, creating new and numerous metabolically active hyphal tips, and extend these to grow into regions of food sources. At the same time, hyphal growth can be considered a form of energy storage, as fungi commonly build up and break down cell-wall material, especially beta-glucans, as a long term energy management strategy (Bartnicki-García, 1999; Wessels, 1966). Additionally, fungi regularly auto-digest contents of older hyphae to recycle nutrients for more active regions of their mycelia (Pollack et al., 2009; White et al., 2002), replacing the organelles of these older hyphal largely with vacuoles, rendering them almost metabolically inert. A lack of true chemotrophy also means that fungi must grow "blindly" to explore their surroundings for new food when substrates are consumed entirely, maintaining approximately the same base level of metabolism per unit mass of active hyphae even while there is no outside energy source available. All of these patterns in fungal growth sum to a very complicated, dynamic picture of respiration and growth in fungi, where the two processes are perhaps indistinguishable.

For the sake of simplicity, we do not attempt to model the complex, non-linear patterns of metabolism fungi as they consume substrates, and search out new substrates in times of scarcity. We simply assume that while a fungal agent has substrate available to consume, it satisfies its energetic and material needs for maintenance of existing biomass, and also harvests a surplus energy unit. When a wood agent is not available to decompose for energy, the fungal agent loses an energy unit per time step, to model the processes of autophagy and scouting growth.

#### Tree placement

Initial tree agent placement on the model landscape follows a "Thomas" process (Thomas 1949), controlled by three, user-defined parameters: the poisson-process rate of parent points that will become centers of tree agent clusters ("kappa" or κ), a secondary gaussian distribution for child points that will become Tree agents ("mu" or μ) the spread (variance) of child points("sigma" or **σ**). Default settings are intended to create approximately 600 tree agents per hectare (Crowther et al 2015). See supplementary jupyter notebook for full details on tree agent placement algorithms.

##### Tree removal

Tree removal can be programmed into model runs at any time. Two types of tree agent removal have been included as functions in the model, to emulate two broad categories of deforestation: (1) thinning, or selective logging, where tree agents are removed at +/- the same rate, throughout the landscape, interspersed among leave tree agents, or (2) fragmenting, where contiguous blocks of forest are removed. The first attempts to emulate the results of selective logging, often in the form of "highgrading." The second is intended to model land use conversions - homesteading, conversion to agriculture, etc. (Kettle and Koh 2014).

Thinning of tree agents requires one argument from the user, the intensity of the thin. This number is between 0 and 1, indicating the proportion of tree agents to be removed, each of which are randomly, independently selected from the pool of the entire set of tree agents on the landscape.

Fragmentation of forest accepts two arguments, the number and radius of fragments. Fragment center locations are assigned randomly, then all tree agents within the user-assigned radius from each center are protected, and the remaining tree agents are removed from the model (fig 4).

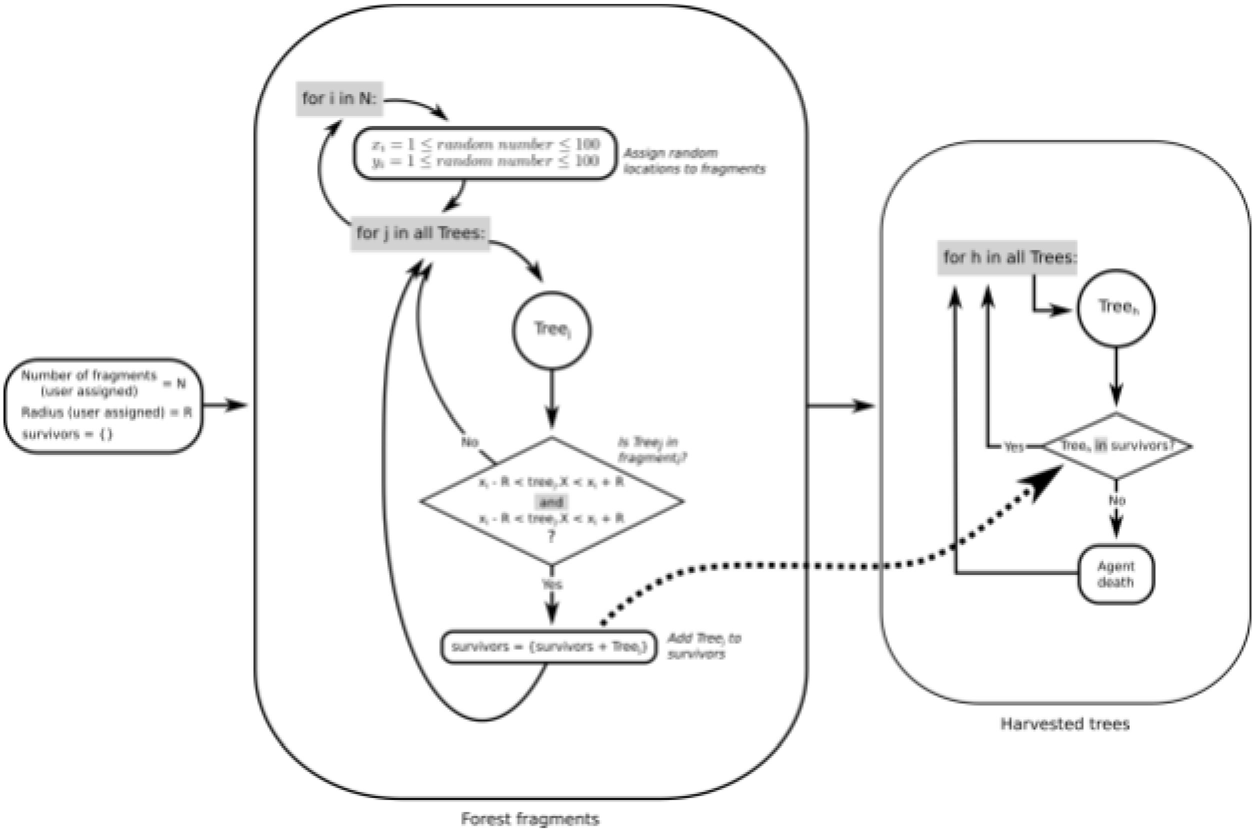


Figure 4. Forest fragmentation submodel. For a higher resolution image click here.
